# Role of endothelial microRNA-150 in pulmonary arterial hypertension

**DOI:** 10.1101/2020.03.25.007021

**Authors:** Giusy Russomanno, Kyeong Beom Jo, Vahitha B. Abdul-Salam, Claire Morgan, Mai Alzaydi, Martin R. Wilkins, Beata Wojciak-Stothard

**Author notes:** Correspondence: Beata Wojciak-Stothard, National Heart and Lung Institute (NHLI), ICTEM Building, Hammersmith Campus, Imperial College London, Du Cane Road, London W12 0NN.

## Abstract

Endothelial dysfunction contributes to the vascular pathology in pulmonary arterial hypertension (PAH). Circulating levels of endothelial miR-150 are reduced in PAH and act as an independent predictor of patient survival. The role of endothelial miR-150 in vascular dysfunction in PAH is not well understood.

Endothelium-targeted miR-150 delivery prevented the disease in Sugen/hypoxia mice, while endothelial knockdown of miR-150 had adverse effects. miR-150 target genes revealed significant associations with PAH pathways, including proliferation, inflammation and phospholipid signaling, with PTEN-like mitochondrial phosphatase (PTPMT1) most markedly altered. PTPMT1 reduced inflammation, apoptosis and improved mitochondrial function in human pulmonary endothelial cells and blood-derived endothelial colony forming cells (ECFCs) from idiopathic PAH. Beneficial effects of miR-150 *in vitro* and *in vivo* were linked with PTPMT1-dependent biosynthesis of mitochondrial phospholipid cardiolipin and reduced expression of pro-apoptotic, pro-inflammatory and pro-fibrotic genes, including *c-MYB, NOTCH3, TGF-β* and *Col1a1*.

In conclusion, we are first to show that miR-150-PTPMT1-cardiolipin pathway attenuates pulmonary endothelial damage induced by vascular stresses and may be considered as a potential therapeutic strategy in PAH.

## INTRODUCTION

Pulmonary arterial hypertension (PAH) is a complex pulmonary vascular disease characterised by excessive vascular endothelial and smooth muscle cell proliferation, inflammation and fibrosis, resulting in increased pulmonary vascular resistance and right ventricular hypertrophy (Schermuly, Ghofrani et al., 2011). The converging effects of hypoxia, inflammation, oxidative stress and metabolic dysregulation contribute to the disease pathology (Bertero, Handen et al., 2018).

At the cellular level, the arterial and right ventricular remodelling in PAH is associated with a shift from oxidative phosphorylation to glycolysis (Freund-Michel, Khoyrattee et al., 2014), which increases the availability of non-oxidized lipids, amino-acids and sugars essential for rapid cell proliferation (Paulin & Michelakis, 2014). These changes are accompanied by inhibition of mitochondrial biogenesis, mitochondrial fragmentation, membrane hyperpolarization and altered ROS production (Culley & Chan, 2018, Paulin & Michelakis, 2014).

Micro RNAs (miRNAs) have emerged as essential regulators of multiple cellular processes, simultaneously controlling mRNA processing, stability and translation of multiple gene targets. Given the multifaceted nature of PAH pathology, there is interest in the role of miRNAs in the pathogenesis of this condition (Negi & Chan, 2017).

We have previously shown that reduced miR-150 levels in plasma, circulating microvesicles and blood cell fraction from PAH patients is a significant predictor of survival, independent of age, cardiac index, disease duration and circulating lymphocyte count (Rhodes, Wharton et al., 2013). A reduction in miR-150 levels in the lung and right ventricle of monocrotaline rats has been shown to correlate with disease severity (Gubrij, Pangle et al., 2016). While miR-150 is highly expressed in mature lymphocytes (Monticelli, Ansel et al., 2005), circulating lymphocytes account for only about 6% of the variation in miR-150 level (Rhodes et al., 2013), suggesting that endothelial cells are a likely source of this miRNA. The impact of variation in endothelial miR-150 expression on endothelial function or disease pathology has not been investigated.

Here we describe the effects of endothelial miR-150 supplementation and inhibition in experimental PAH, human pulmonary artery endothelial and smooth muscle cells and blood-derived endothelial colony forming cells (ECFCs) from PAH patients and identify the signaling mediators involved. We show that miR-150 has anti-apoptotic, anti-inflammatory, anti-proliferative and anti-fibrotic effects and is required for mitochondrial adaptation to increased energy demand in conditions of vascular stress.

## RESULTS

### Endothelial supplementation of miR-150 improves pulmonary vascular haemodynamics and reduces vascular remodelling in Sugen/hypoxia mice

The effect of miR-150 administration was evaluated in the Sugen/hypoxia mouse, an experimental model of pulmonary hypertension (Fig 1A). miR-150-5p complexed with a lipid carrier DACC was delivered at 4-day intervals throughout the three week period of study. The DACC liposomal formulation targets the vascular endothelium, with highest efficiency seen in the lung (Abdul-Salam, Russomanno et al., 2019, Fehring, Schaeper et al., 2014). The control Sugen/hypoxia animals (treated with scrambled miR control) showed significantly elevated RVSP, right ventricular hypertrophy (RVH) and increased intrapulmonary small vessel muscularisation marked by a prominent staining of α-SMA of pre-capillary arterioles (Figure 1 B-E). Lung and heart levels of miR-150-5p were significantly reduced in these mice when compared with healthy transfection controls (Figure 1F, G). *In situ* hybridization analysis showed that the most prominent reduction in miR-150 expression was in the vascular endothelium (Appendix Figure S1), while miR-150 expression levels in leukocytes and airway epithelium remained relatively unaffected.

**Figure 1.**
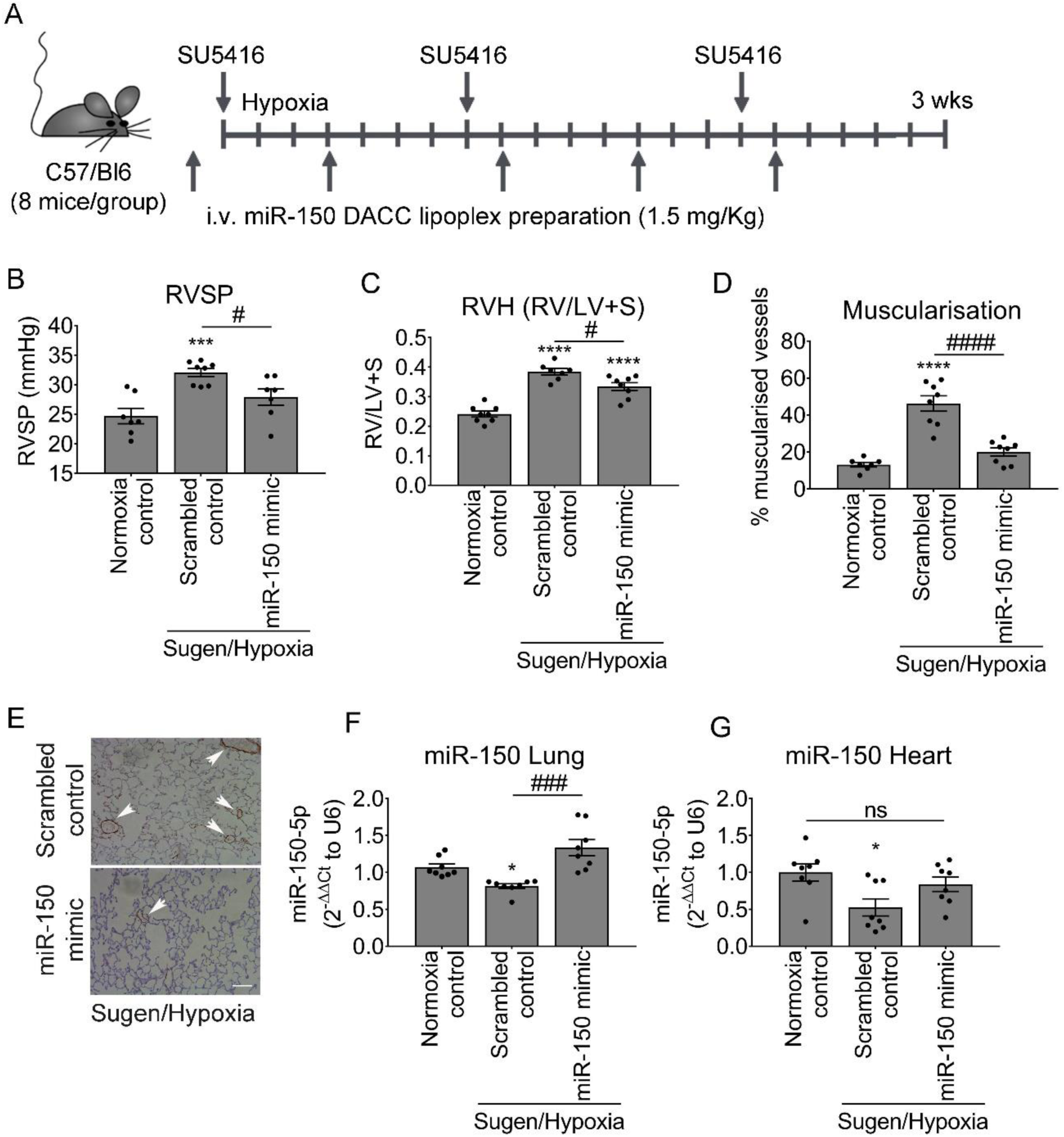
Effect of pulmonary endothelial miR-150 supplementation on development of PH in Sugen/hypoxia mice. (A) Experimental layout. (B) Right ventricular systolic pressure (RVSP), (C) Right ventricular hypertrophy (RVH: right ventricle to left ventricle + septum ratio RV/LV+S), (D) percentage of muscularised vessels <50 µm in diameter/total number of vessels in the lungs of normoxia control mice and Sugen/hypoxia mice treated with scrambled control or miR-150 mimic delivered by iv administration of DACC lipoplex, as indicated. (E) Representative images of αSMA staining in lung sections from mice treated with scrambled control or miR-150 mimic. (F, G) miR-150 expression in lung and heart, as indicated. *p<0.05, **p<0.005, ***p<0.001, ****p<0.0001, comparisons with normoxia control; ^#^p<0.05, ^###^p<0.001, ^####^p<0.0001, comparisons with scrambled control, Bars are means ± SEM; N=8 mice/group, one-way ANOVA with Tukey post-test.

Endothelium-targeted DACC/miR-150 delivery increased expression levels of miR-150 in lung and heart tissues in Sugen/hypoxia mice compared with controls (Figure 1F, G). The treatment reduced RVSP (p<0.05), RVH (p<0.05) and vascular muscularisation (p<0.0001; Figure 1B-E).

### Heterozygous endothelial-specific deletion of miR-150 worsens the symptoms of PH

To evaluate the effects of endothelium-specific depletion of miR-150, mice with inducible conditional endothelium-specific deletion of miR-150 were obtained by crossing floxed miR-150 mice with mice carrying tamoxifen-inducible Cre recombinase under the control of the Cdh5 promoter (Cdh5(PAC)-iCreERT2) (Birdsey, Shah et al., 2015). Only heterozygous miR-150iEC-KO mice (miR-150^fl^/Cdh5(PAC)-iCreERT2) were used for experiments to mimic the reduction (but not complete depletion) of miR-150 content seen in human disease and pre-clinical models of PAH. Following tamoxifen administration, the efficiency of Cre-recombinase deletion of miR-150 was confirmed by qPCR (Figure 2A).

**Figure 2.**
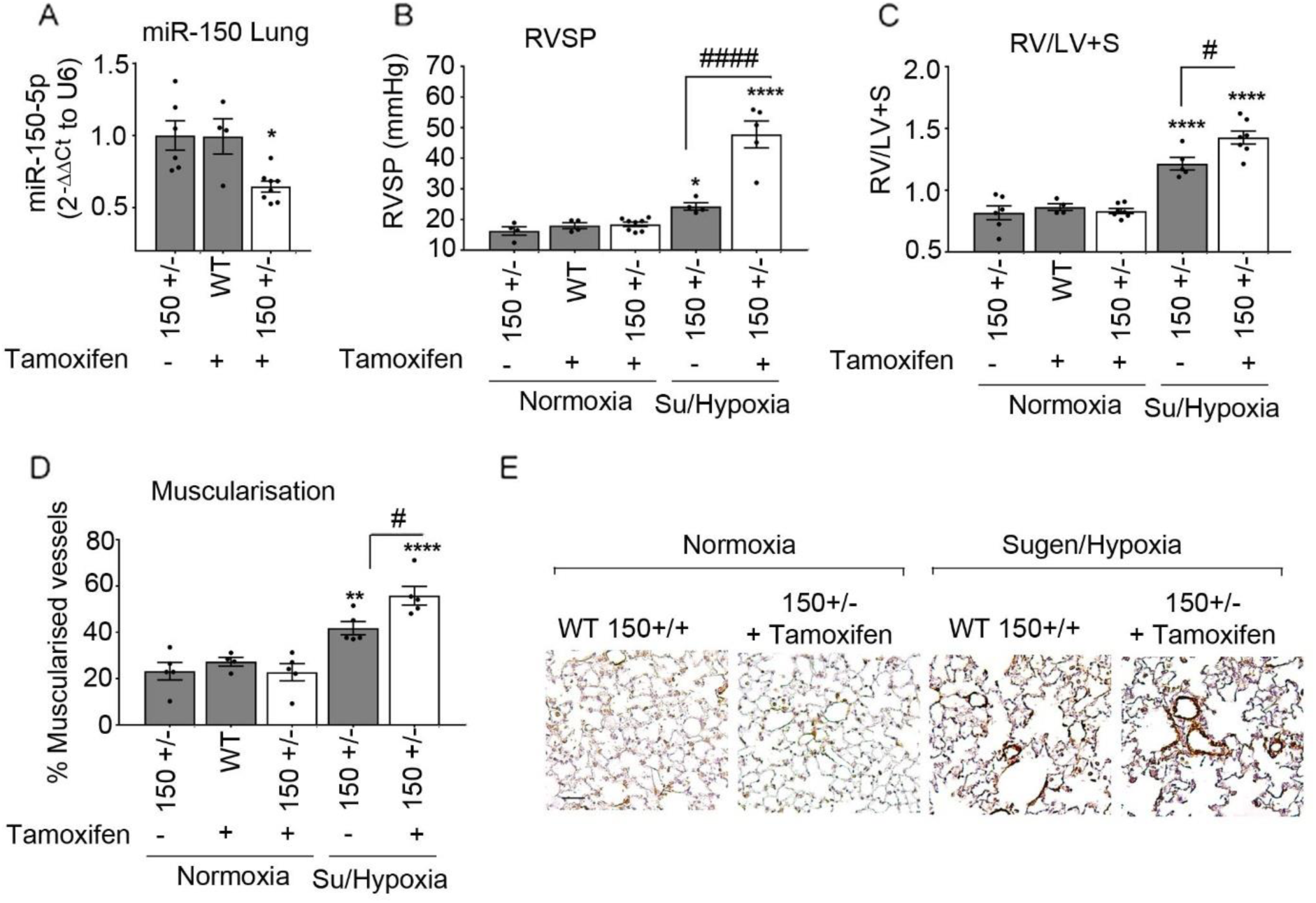
Effect of endothelial miR-150 deletion on development of PH in Sugen/hypoxia mice. (A) Effect of tamoxifen administration on miR-150 levels in lungs of WT (150^+/+^) and miR-150iEC-HTKO mice (150^+/−^). (B) RVSP, (C) RV/LV+S, (D) percentage of muscularised vessels <50 µm in diameter/total number of vessels in the lungs of normoxia wildtype control (+ tamoxifen), normoxia miR-150^+/−^ control (without tamoxifen) and Sugen/Hypoxia miR-150^+/−^ mice with and without tamoxifen, as indicated. In (A-D) empty bars mark miR-150-deficient animals. (E) Representative images of α-SMA staining. Bar=25µm *p<0.05, **p<0.005, ****p<0.0001, comparisons with normoxia control; ^#^p<0.05, ^####^p<0.0001, comparisons with miR-150^+/−^ control. Bars are means ± SEM; N=4-8 mice/group, one-way ANOVA with Tukey post-test.

Sugen/hypoxia miR-150iEC-KO mice showed a substantial elevation of RVSP (∼2-fold increase, p<0.0001), accompanied by rise in RVH and pulmonary vascular muscularization (both p<0.05), compared with Sugen/hypoxia wildtype littermates (Figure 2 B-E).

### Identification of miR-150-regulated genes

In order to identify potential mediators of miR-150-induced effects, HPAECs transfected with miR-150 or non-targeting control miRNA were subjected to RNA profiling.

Out of the 13,767 genes identified, 180 genes were significantly upregulated (p<0.01, fold change>1.5) and 207 downregulated (p<0.01, fold change<-1.5) by miR-150 (Figure 3A). Heat map and unsupervised hierarchical clustering of the top 26 differentially expressed genes (with adjusted p-value<0.05) are shown in Figure 3B. A list of differentially expressed genes is provided in Appendix Table S1.

**Figure 3.**
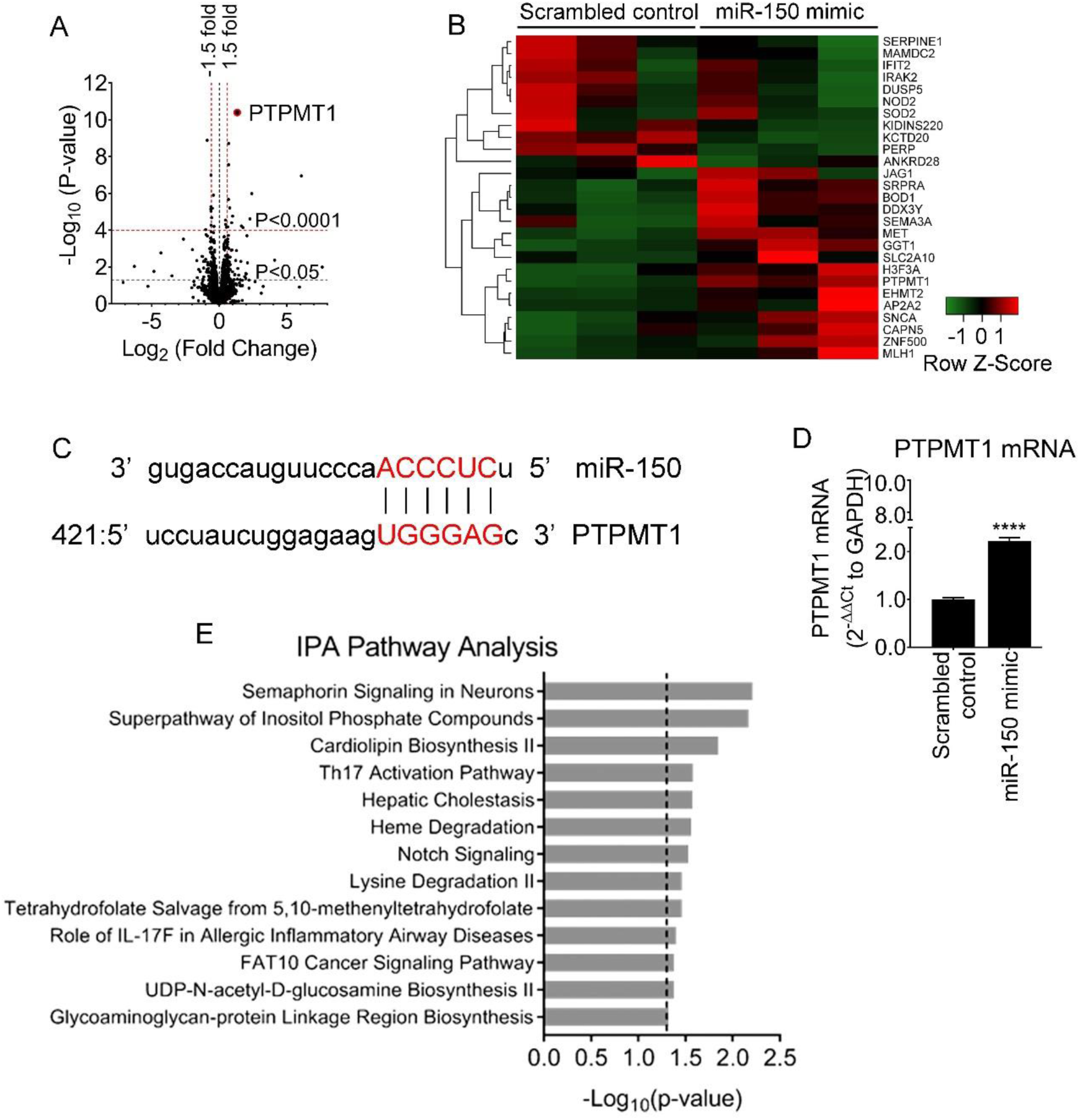
RNA-sequencing analysis in miR-150 overexpressing HPAECs shows PTPMT1 as the most upregulated gene. To identify miR-150 signaling mediators, HPAECs were transfected withmiR-150-5p or scrambled control (20 nM) and RNA was extracted for RNA-sequencing (in three independent experiments). (A)Volcano plot of differentially expressed genes (DEG). Each point represents the difference in expression (Log2 fold difference) between groups and the associated significance of this change (independent unpaired sample t-test). PTPMT1 is highlighted as the most up-regulated gene (fold change=2.51, p=3.98×10^−11^, adjusted p-value=5.48×10^−7^; N=3). (B) Heat map showing 26 most significant genes after multiple test correction using the Benjamini-Hochberg procedure. Green and red represent downregulation and upregulation, respectively. (C) PTPMT1 mRNA levels in cells transfected with control miRNA and miR-150-5p. (D) miR-150 binding sequence in the promoter region of PTPMT1. (E) IPA pathway enrichment analysis in cells transfected with miR-150.

PTPMT1, a mitochondrial protein tyrosine phosphatase essential for cardiolipin biosynthesis, was the most significantly altered gene, showing a 2.5-fold increase in expression (adjusted p-value=5.48×10^−7^) compared with controls. miRNAs can reduce gene expression by binding to the 3’UTR of target mRNAs or increase target gene expression by binding to gene promoters(Vaschetto, 2018). miR-150-5p binding sequences are present in the *PTPMT1* promoter (Fig. 3C), likely to indicate a direct transcriptional regulatory effect. Increase in the expression of *PTPMT1* and reduction in the expression of other gene regulators of vascular remodelling identified by RNAseq, *SERPINE1* (Simone, Higgins et al., 2014), *PERP* (Zhao, Chen et al., 2019), *DUSP5* (Chen, Zmuda et al., 2020*), NOTCH3* (Li, Zhang et al., 2009) and *c-myb* (You, Mungrue et al., 2003) were validated by qPCR (Fig. 3D and Appendix Figure S2).

IPA pathway enrichment analysis of miR-150 gene targets showed significant associations with pathways regulating cell proliferation, inflammation and oxidative metabolism, including NOTCH signaling (p=0.029), cardiolipin biosynthesis (p=0.014) and inositiol phosphate signaling (p=0.007) (Fig. 3E).

Consistent with the findings *in vitro*, quantification of transcripts by qPCR and RNAscope fluorescent *in situ* hybridization, which allows specific identification and quantification of single transcripts (Wang, Flanagan et al., 2012), confirmed upregulation of *PTPMT1* (Fig. 6A) and downregulation of miR-150 target genes, *c-myb* and *NOTCH3*, in the lungs of miR-150-treated animals (Fig. 6A and Appendix Figures S3 and S4). Heart tissue from miR-150-treated Sugen/hypoxia mice showed significantly reduced expression of markers of cardiac hypertrophy and fibrosis, including transforming growth factor beta (TGF-β), alpha-1 type I collagen (Col1a1) and regulator of calcineurin 1 (Rcan1) (Appendix Figure S5). In contrast, heart tissue from Sugen/hypoxia miR-150iEC-KO mice showed significantly increased levels of Col1a1, compared with the corresponding wildtype disease controls (Appendix Figure S6).

**Figure 4.**
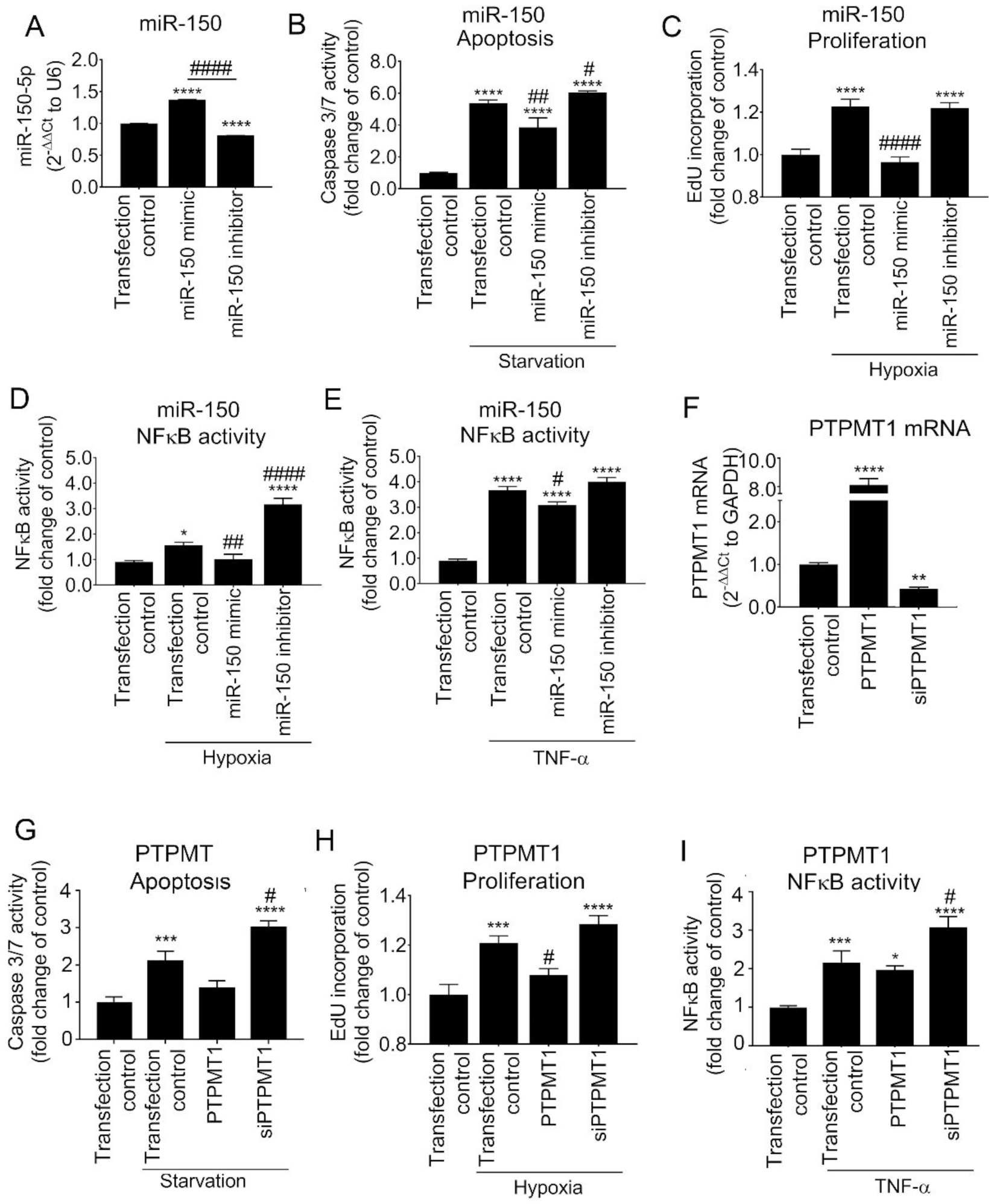
Endothelium-protective effects of miR-150 and PTPMT1. (A-E) Effect of miR-150 mimic and miR-150 inhibitor on (A) miR-150 expression levels in HPAECs, (B) starvation-induced apoptosis (caspase 3/7 activity assay), (C) hypoxia-induced proliferation (EdU incorporation assay), (D) hypoxia (24h)- and (E) TNF-α-induced (10 nmol/L, 24h) NFκB activity (luciferase reporter assay). (F-I) Effect of PTPMT1 or overexpression or silencing (siPTPMT1) on (F) PTPMT1 mRNA expression, (G) apoptosis, (H) proliferation and (I) TNF-α-induced NFkB activity in HPAECs. In (A) N=3, in (B-I) N=6. *p<0.05, **p<0.005, ***p<0.001, ****p<0.0001, comparisons with untreated transfection control; ^#^p<0.05, ^##^p<0.001, ^####^p<0.0001, comparisons with treated transfection control. Bars are means ± SEM; one-way ANOVA with Tukey post-test.

**Figure 5.**
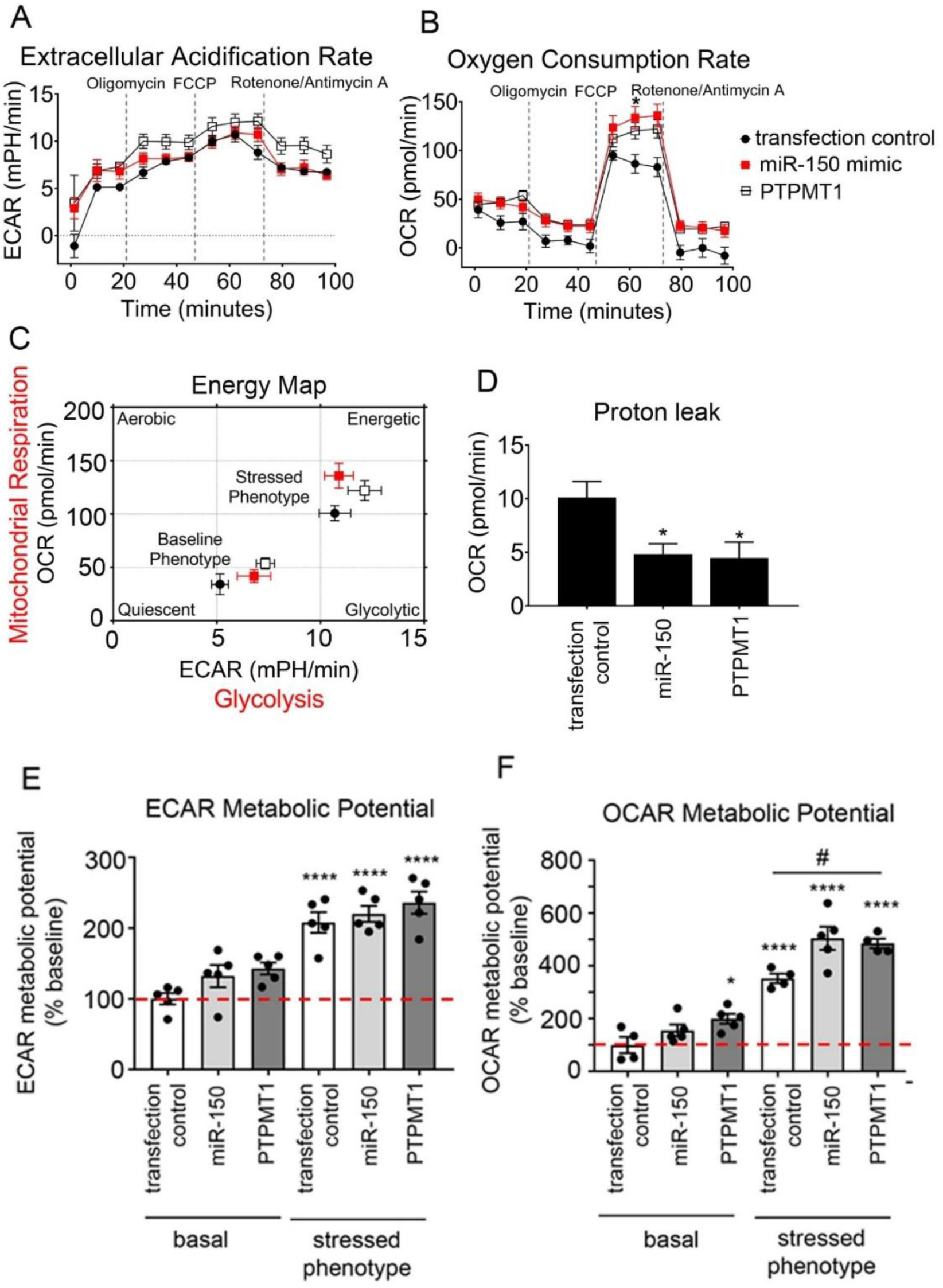
Effect of miR-150 and PTPMT1 on metabolic potential in HPAECs. (A) Extracellular acidification rate (ECAR), (B) oxygen consumption rate (OCR), (C) energy map, (D) proton leak, (E) ECAR metabolic potential (% of basal control), and (F) OCR metabolic potential (% of basal control) in control HPAECs (transfection control) and HPAECs transfected with miR-150 or PTPMT1, as indicated. N=5. Bars are means ± SEM; one-way ANOVA with Tukey post-test. *p<0.05, ****p<0.0001, comparison with transfection control (basal). ^#^p<0.05, as indicated.

**Figure 6.**
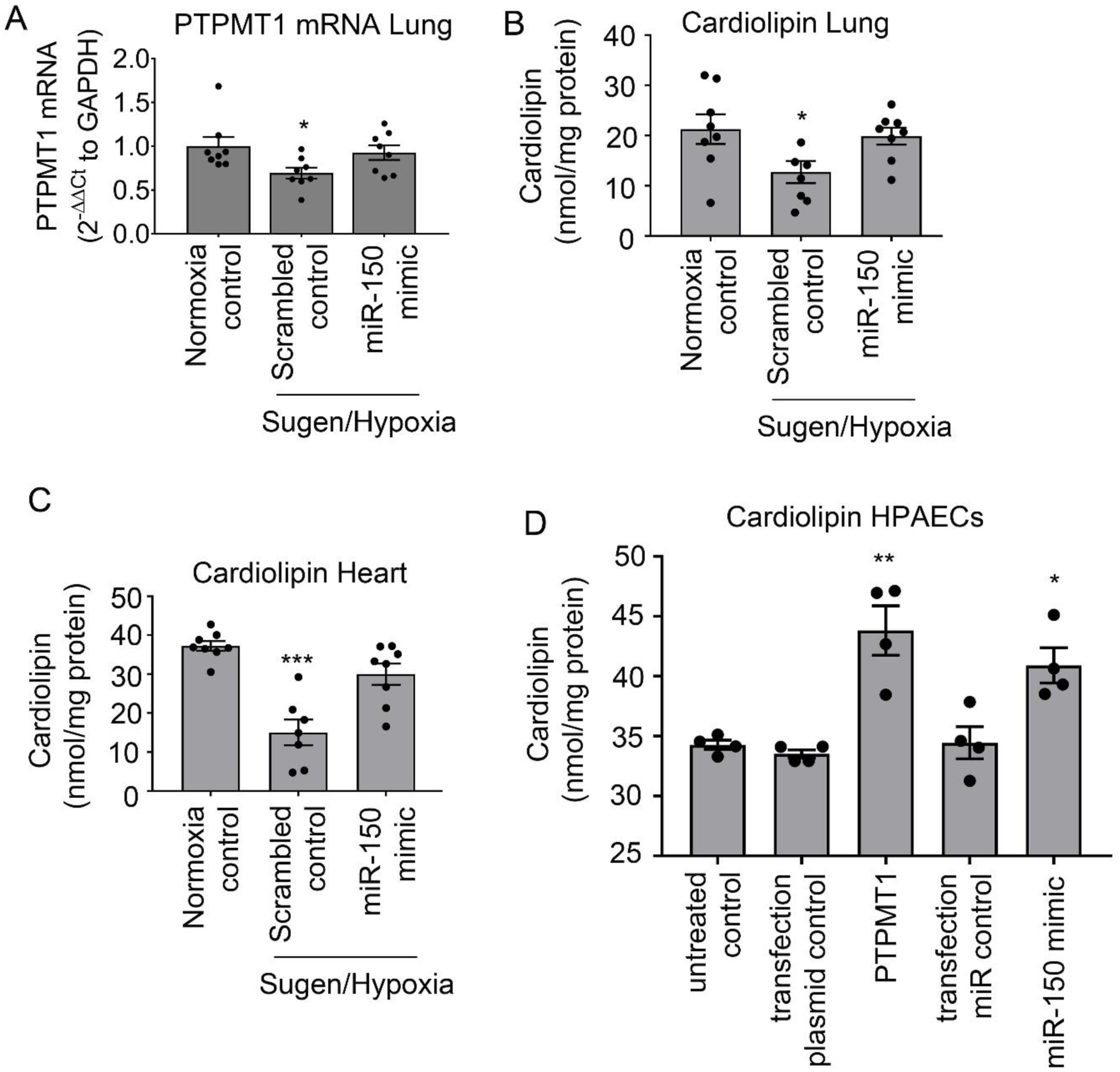
miR-150 and PTPMT1 modulate cardiolipin levels in tissues and cells. (A) Lung PTPMT1, (B) lung cardiolipin, (C) heart cardiolipin levels in control and DACC/miR-150-treated Sugen/hypoxia mice, as indicated. (D) Cardiolipin levels in control HPAECs and HPAECs transfected with miR-150 and PTPMT1, as indicated. In graphs, bars are means ± SEM; one-way ANOVA with Tukey post-test. *p<0.05, **p<0.01, ***p<0.001, comparison with transfection control. N=4-8.

### PTPMT1 mediates homeostatic effects of miR-150

Transfection of HPAECs with miR-150 markedly attenuated endothelial cell apoptosis, hypoxia-induced cell proliferation and hypoxia- and TNF-α-induced NFκB activation (Figure 4A-E). In contrast, inhibition of miR-150 markedly augmented endothelial damage and inflammation (Figure 4B-E). Transfection efficiency evaluated with Cy5-labelled miR negative control was∼ 85% (Appendix Figure S7). Changes in the intracellular miR-150 levels in cells transfected with miR-150 mimic and inhibitor were confirmed by qPCR (Fig. 4A).

To see whether manipulation of miR-150 levels in endothelial cells can affect smooth muscle cell proliferation, HPAECs and HPASMCs were seeded on the opposite sides of a porous membrane in Transwell dishes (Appendix Figure S8). Endothelial miR-150 overexpression significantly reduced hypoxia-induced proliferation of HPASMCs (Appendix Figure S8)

Overexpression and silencing of *PTPMT1* mimicked to large extent changes induced by miR-150 mimic and miR-150 inhibitor, respectively (Figure 4 F-I), suggesting that PTPMT1 acts as a key mediator of the anti-proliferative and anti-inflammatory effects of miR-150 in pulmonary endothelial cells.

### miR-150 and PTPMT1 improve mitochondrial function in HPAECs

Energy metabolism constitutes an essential link between cell growth and apoptosis (Mason & Rathmell, 2011). In order to assess the effect of miR-150 and PTPMT1 on energy metabolism, HPAECs and HPAECs transfected with miR-150 or PTPMT1 were subjected to bioenergetic profiling. Extracellular acidification rate (ECAR), which reflects the level of glycolysis, was not significantly affected by either treatment, but mitochondrial oxygen consumption rate (OCAR), reflective of the level of mitochondrial respiration, was significantly elevated in miR-150 and PTPMT1-overexpressing cells (Figure 5A-C).

The treatment of cells with miR-150 and PTPMT1 significantly reduced mitochondrial proton leak (Figure 5D). As proton leak depicts the protons that migrate into the matrix without producing ATP, a reduction in proton leak indicates an improvement in coupling of substrate oxygen and ATP generation (Cheng, Nanayakkara et al., 2017).

Measurement of metabolic potential helps to evaluate the capacity of cells to respond to stress conditions associated with increased energy demand via mitochondrial respiration and glycolysis. The results show that miR-150 and PTPMT1 significantly increased mitochondrial metabolic potential measured as fold-increase in OCAR over basal control levels, while glycolytic metabolic potential in cells remained relatively unaffected (Figure 5E, F).

### miR-150 and PTPMT1 restore cardiolipin levels in Sugen/hypoxia lung and heart tissues and increase mitochondrial content in human PAH ECFCs

PTPMT1 is a mitochondrial tyrosine kinase, essential for the biosynthesis of cardiolipin, the main phospholipid component of mitochondrial membranes and a key regulator of mitochondrial structure and function (Dudek, 2017, Shen, Liu et al., 2011). We evaluated the effect of miR-150 and PTPMT1 supplementation on cardiolipin levels in lungs and hearts from miR-150-treated Sugen/hypoxia mice, HPAECs and ECFCs from IPAH patients.

PTPMT1 and cardiolipin levels were significantly reduced in Sugen/hypoxia mice, while miR-150 supplementation restored their expression to the level seen in healthy mice (Figure 6A-C). Overexpression of PTPMT1 and miR-150 significantly elevated cardiolipin levels in cultured endothelial cells (p<0.01 and p<0.05, respectively) (Figure 6D).

Blood-derived endothelial colony-forming cells (ECFCs) are often used as surrogates for pulmonary endothelial cells in PAH(Duong, Comhair et al., 2011). qPCR analysis showed that miR-150 and PTPMT1 expressions were markedly reduced in ECFCs from IPAH patients, compared with the cells from healthy individuals (p<0.01, n=12-14) (Figure 7 A, B). IPAH cells also showed a marked (∼2-fold, p<0.05) reduction in cardiolipin levels, which was restored upon treatment with miR-150 and PTPMT1 (Figure 7C).

**Figure 7.**
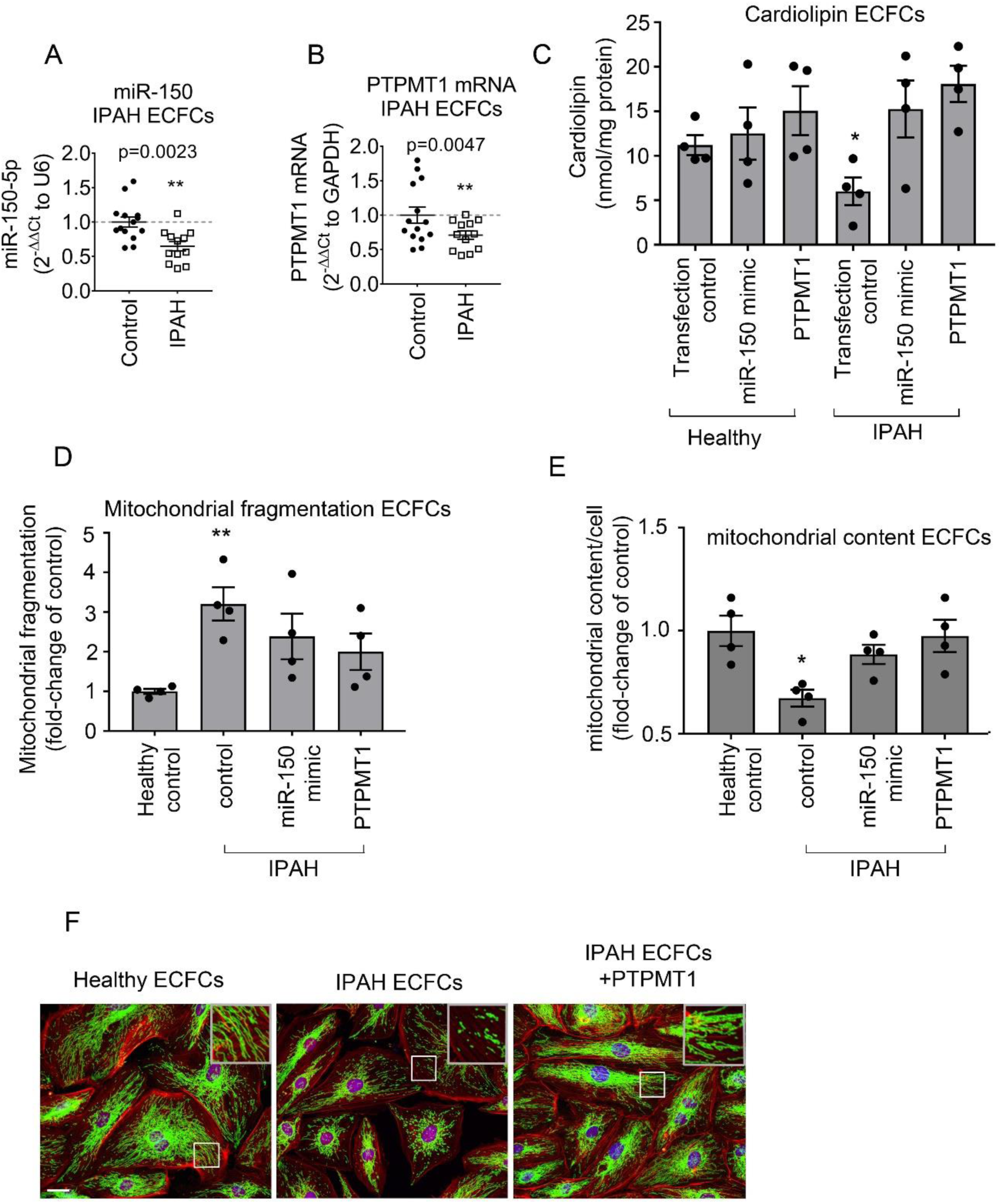
miR-150, PTPMT1 and cardiolipin levels and mitochondrial biogenesis in IPAH ECFCs. (A) miR-150 and (B) PTPMT1 expression levels in ECFCs from healthy individuals and IPAH patients; (C) Cardiolipin levels; (D) mitochondrial fragmentation and (E) mitochondrial content (mitochondrial coverage/cell) in ECFCs treated, as indicated. (F) representative images of mitochondrial fragmentation in healthy and IPAH ECFCs. Inset in the top right corner is an enlarged image of the boxed area. Mitochondria were immunolabelled with FITC (green) and F-actin with TRITC-phalloidin (red). Bar= 10 µm. Data are expressed as means ± SEM *p<0.05, **p<0.01, comparison with healthy control. In (A, B) N=12-14 and in (C-E) N=4.

Reduction in mitochondrial oxidative phosphorylation in PAH is linked with an increase in mitochondrial fragmentation and a reduction in mitochondrial biomass(Culley & Chan, 2018). Accordingly, we observed an increased mitochondrial fragmentation and reduction in mitochondrial content in ECFCs from IPAH patients compared with healthy controls (Figure 7 D-F). These changes were reversed by overexpression of miR-150 and PTPMT1 (Figure 7 D-F).

## DISCUSSION

This study is first to demonstrate the vasculoprotective effects of endothelial miR-150 in PAH. Our data show that miR-150 facilitates vascular adaptation to stress conditions by increasing mitochondrial metabolic potential via increased expression of PTPMT1, the key regulator of cardiolipin biosynthesis. The vasculoprotective effects of endothelial miR-150 are associated with reduced expression of markers of inflammation, apoptosis and fibrosis critical to the pathology of PAH, including c-myb, NOTCH3, TGF-β and Col1a1. A schematic diagram of miR-150-induced effects is shown in Figure 8.

**Figure 8.**
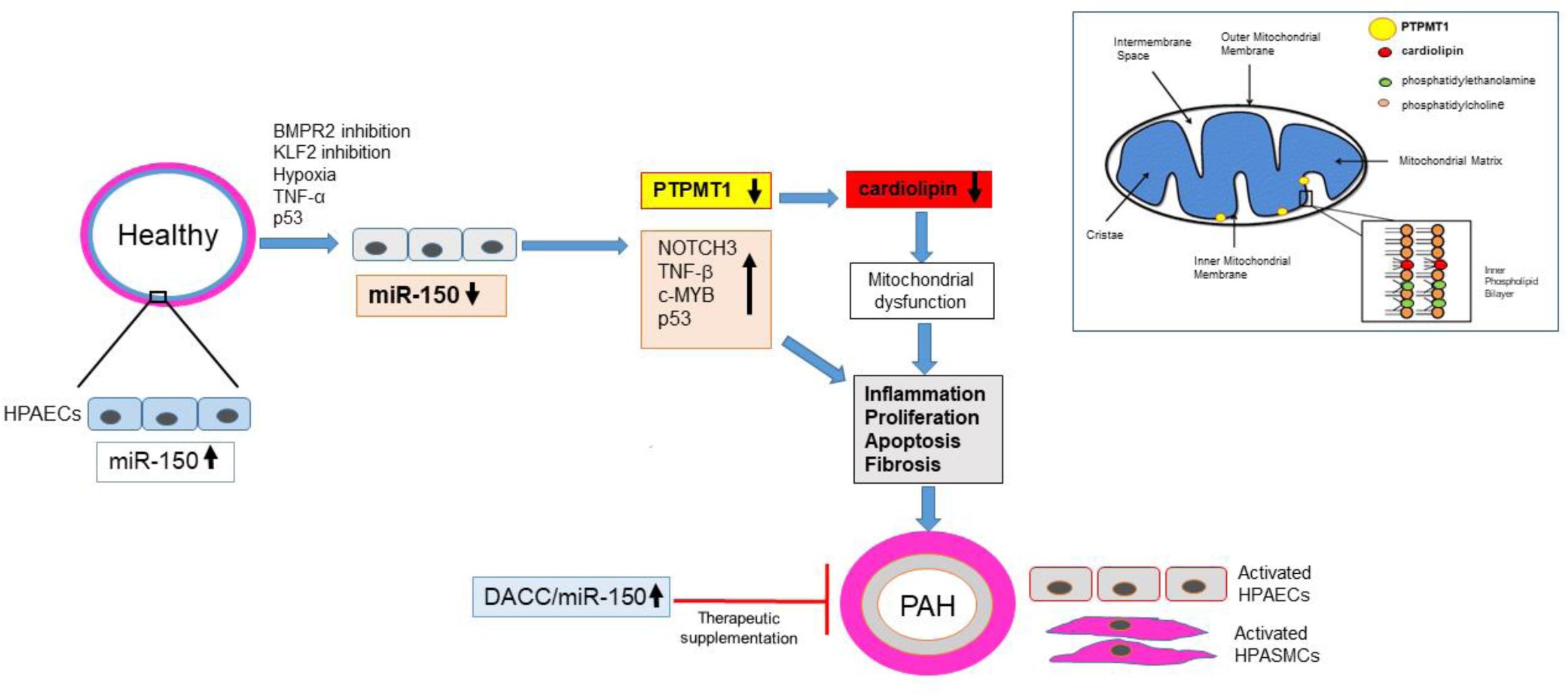
Schematic diagram of the proposed miR-150 signaling pathway. The inset shows localization of PTPMT1 and cardiolipin within the mitochondrion.

Downregulation of miR-150 in pulmonary endothelium and IPAH endothelial cells is likely to result from transcriptional repression by apoptosis-inducer p53 (Ghose & Bhattacharyya, 2015), highly expressed in endothelial cells in pulmonary hypertensive lung (Wang, Yang et al., 2019). An alternative mechanism may involve inhibition of flow-activated transcription Krueppel-like factor 2 (KLF-2), a known inducer of miR-150 expression (Hergenreider, Heydt et al., 2012). Recent studies show that KLF2 signaling is compromised and expression levels of KLF2-induced miRNAs are reduced in PAH (Chandra, Razavi et al., 2011, Eichstaedt, Song et al., 2017, Sindi, Russomanno et al., 2020). Hypoxia (Chen, Shen et al., 2017) and inhibition of bone morphogenetic protein receptor 2 (BMPR2) (Eichstaedt et al., 2017) are likely to play a contributory role.

RNA sequencing identified *PTPMT1* as a key gene affected by miR-150 overexpression and functional analysis showed that PTPMT1 is a key mediator of endothelium-protective effects of miR-150. PTPMT1 is exclusively localized to the inner membrane of mitochondria, with close proximity to electron transport chain complexes and enzymes of the tricarboxylic acid cycle (Shen et al., 2011). Interestingly, PTPMT1-ablated cells show a marked decrease in aerobic metabolism and enhancement of glycolysis (Shen et al., 2011), reminiscent of changes seen in PAH. In addition, inhibition of PTPMT1 is known to increase mitochondrial fragmentation (Zhang, Guan et al., 2011), a hallmark of PAH (Dasgupta, Wu et al., 2020).

ECFCs from PAH patients show metabolic reprogramming similar to the one seen in PAH lung (Archer, 2017, Caruso, Dunmore et al., 2017), providing a rationale for the use of these cells in disease modeling. Consistent with the pathology observed in pulmonary endothelial and smooth muscle cells from PAH lungs (Dasgupta et al., 2020), PAH ECFCs showed reduced mitochondrial content and increased mitochondrial fragmentation, which were prevented by overexpression of miR-150 and PTPMT1.

The effects of PTPMT1 can be linked to its role in the biosynthesis of cardiolipin, a mitochondrial-specific phospholipid regulating mitochondrial membrane integrity and function. PTPMT1 dephosphorylates phosphatidylglycerophosphate, resulting in an increase in phosphatidylglycerol, an essential intermediate for biosynthesis of cardiolipin. Interaction with cardiolipin is required for optimal activity of several inner mitochondrial membrane proteins, including the enzyme complexes of the electron transport chain and ATP production. Moreover, cardiolipin plays an important role in mitochondrial membrane morphology, stability and dynamics, in mitochondrial biogenesis and mitochondrial fusion (Dudek, 2017).

Our data argue for a vasculo- and cardio-protective role for the miR-150-PTPMT1-cardiolipin pathway in PAH. Inhibition of cell apoptosis by miR-150 can be linked with attenuation of TNF-α-induced mitochondrial dysfunction by PTPMT1 and cardiolipin (Choi, Gonzalvez et al., 2007) or reduced expression of apoptosis markers, p53 and EGR2 (Liu & Di Wang, 2019, Wu, Jin et al., 2010).

miR-150 and PTPMT1 may reduce maladaptive RV remodeling by augmentation of glucose oxidation and prevention of capillary rarefaction (Ryan & Archer, 2014). miR-150 has been shown to increase arteriolar density and improve blood flow in ischaemic heart tissues in mouse models of hypercholesterolemia (Desjarlais, Dussault et al., 2017). Consistently, we observed that miR-150 reduced expression of markers of cardiac hypertrophy and fibrosis, TGF-β, Col1a1, Rcan1 (Voelkel, Quaife et al., 2006) in Sugen/hypoxia mice.

Endothelial-cardiomyocyte communication is an key regulator of cardiac function (Lim, Lam et al., 2015). Increased ROS generation linked to endothelial apoptosis in pre-clinical models of PAH, contributes to RV failure (Paradies, Paradies et al., 2019, Rawat, Alzoubi et al., 2014). Electron leak is the major causative factor for production of mitochondrial superoxide, hence the reduction in mitochondrial proton leak by miR-150 and PTPMT1 may account for the beneficial effects of miR-150 treatment. Interestingly, low cardiolipin levels have been linked with reduced mitochondrial complex II + III activity and RV failure in MCT rats and porcine model of persistent pulmonary hypertension of the newborn (Saini-Chohan, Dakshinamurti et al., 2011).

Anti-remodeling effects of miR-150 are likely to result from the cumulative changes in expression of multiple genes and delineating individual contributions of miR-150 targets is beyond the scope of this investigation. Besides PTPMT1, other signaling mediators, including c-MYB, NOTCH3, activin receptors 1 and 2 and matrix metalloproteinases, are likely to play a role (Cao, Hou et al., 2014, Rhodes et al., 2013). c-MYB stimulates cell migration (Li, Zhang et al., 2013), increases recruitment of endothelial progenitor cells (Wang, Li et al., 2014) and promotes cardiac hypertrophy and fibrosis (Deng, Chen et al., 2016). NOTCH3 is a marker and predictor of PAH (Boucher, Gridley et al., 2012, Li et al., 2009) and its blockade is sufficient to reverse experimental PAH (Havrda, Johnson et al., 2006, Li et al., 2009). Consistently, we observed contemporaneous, opposing changes in the expression of miR-150 and its targets, c-MYB and NOTCH3, in human cells and lung tissues from PAH mice.

To summarise, we show that reduction in endothelial miR-150 levels has an adverse effect on pulmonary haemodynamics in PAH, while endothelium-targeted delivery of miR-150 is protective. Improvement of mitochondrial function by miR-150-PTPMT1-cardiolipin signaling is likely to facilitate adaptation of lung and heart to high energy demand created by mechanical workload, inflammation and hypoxia in stress conditions.

## MATERIALS AND METHODS

### Animal experiments

All studies were conducted in accordance with UK Home Office Animals (Scientific Procedures) Act 1986 and ARRIVE guidelines. All animals were randomly allocated to groups, and all personnel involved in data collection and analysis (haemodynamics and histopathologic measurements) were blinded to the treatment status of each animal. Only weight- and age-matched males were included for experimentation as, in contrast to the human clinical studies, most animal studies have shown that female sex and estrogen supplementation have a protective effect against PAH.

To induce PAH, 8-12 weeks old C57/BL male mice (20 g; Charles River, UK) were injected subcutaneously with Sugen (SU5416; 20mg/kg), suspended in 0.5% [w/v] carboxymethylcellulose sodium, 0.9% [w/v] sodium chloride, 0.4% [v/v] polysorbate 80, 0.9% [v/v] benzyl alcohol in deionized water once/week. Control mice received only vehicle. Mice were either housed in normal air or placed in a normobaric hypoxic chamber (10% O_2_) for 3 weeks (n= 8/group).

mirVana hsa-miR-150-5p (ID MC10070) mimic or scrambled miRNA control (Ambion) in complex with DACC lipoplex preparation (Silence Therapeutics)(Fehring et al., 2014) was administered intravenously once every fourth day at 1.5 mg/kg/day for 3 weeks, on 5 occasions. The first injection was given 1 day before Sugen/hypoxia administration. At 3 weeks, the mice were anaesthetised by intraperitoneal injection of Ketamine/Dormitor (75 mg/kg + 1 mg/kg).

To produce mice with inducible conditional endothelium-specific deletion of miR-150, floxed miR-150 mice (STOCK Mir150^tm1Mtm^/Mmjax mice from Jackson Laboratories) C57/Bl6 background were crossed with C57/Bl6 mice carrying tamoxifen-inducible Cre recombinase under the control of the Cdh5 promoter (Cdh5(PAC)-iCreERT2 (Wang, Nakayama et al., 2010). Following tamoxifen administration, efficient Cre-recombinase deletion of miR-150 was confirmed by PCR in miR-150^fl/fl^/Cdh5(PAC)-iCreERT2 mice (henceforth referred to as miR-150iEC-KO).

Weaned mice were ear-notched and samples were incubated in lysis buffer (100 mM Tris HCl pH 8.5, 5 mM EDTA, 200 mM NaCl, 0.2% SDS, 0.14 mg/mL Proteinase K, Ambion™) for 2 hours at 55°C under agitation (700 rpm). Samples were then vortexed and pelleted at 14,000 rpm for 10 minutes. Supernatant was transferred to a new DNase-free tube and DNA was precipitated in isopropanol (20 minutes incubation at RT). DNA was pelleted at 14,000 × g for 10 minutes, supernatant was discarded and the DNA pellet was air dried and then resuspended in 100 µL of DNase-free water.

PCR reactions were performed using REDTaq^®^ ReadyMix™ PCR Reaction Mix (Sigma-Aldrich, cat. R2523) with the primers listed in Table 1 (500 nM of each) in a SmplyAmp™ Thermal Cycler (Applied Biosystems). All primers were purchased from Sigma-Aldrich.

**Table 1.**
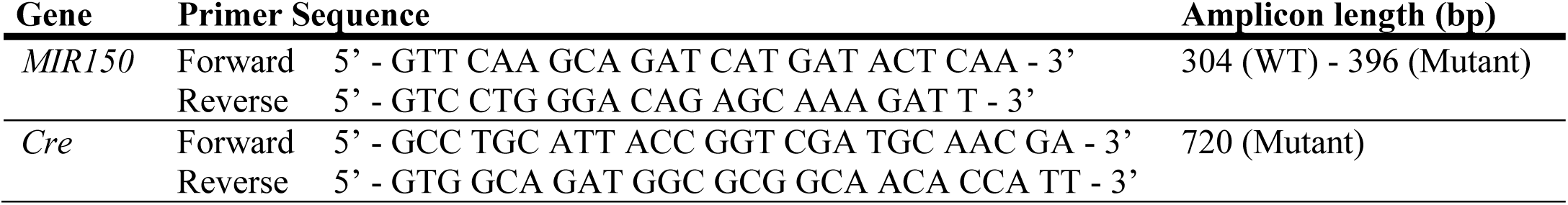
Sequences of specific primers used for mouse genotyping.

Thermocycling conditions for miR-150 followed the Jackson Laboratory’s instructions (www.jax.org): 2 min at 94°C, then 20 s at 94°C, 15 s at 65°C, 10 s at 68°C, for 10 cycles, 15 s at 94°C, 15 s at 60°C, 10 s at 72°C, for 28 cycles, and a last step of 2 min at 72°C. For Cre genotyping, thermocycling conditions were as follows: 3 min at 94°C, then 30 s at 94°C, 30 s at 70°C, 60 s at 72°C, for 32 cycles, and a last step of 10 min at 72°C.

All PCR products were separated on a 2% agarose gel, visualized using GelRed Nucleic Acid Gel Stain (Thermo Fisher Scientific, cat. NC0017761), and size was estimated with comparison to a DNA mass ladder (GeneRuler 100 bp DNA Ladder, Thermo Fisher Scientific, cat. SM0243).

#### Sugen/hypoxia model of PAH

At 6 weeks of age, miR-150^fl^/Cdh5(PAC)-iCreERT2 mice were injected intraperitoneally with 100 µL of 5 mg/mL tamoxifen (Sigma, cat. no. T5648) or vehicle (12.5% vol/vol ethanol in peanut oil) for 5 consecutive days (Pitulescu, Schmidt et al., 2010). Littermate wild-type animals were used as control. Two weeks after tamoxifen injection, mice were injected with Sugen and housed in normal air or hypoxia as previously described for 4 weeks (n=4-8/group).

The development of PAH was verified by measuring right ventricular systolic pressure (RVSP), right ventricular hypertrophy (assessed as the right ventricle to left ventricle/septum ratio -RV/LV+S), and muscularisation of small intrapulmonary arteries, as previously described(Wojciak-Stothard, Abdul-Salam et al., 2014). RVSP was measured via direct cardiac puncture using a closed-chest technique in the spontaneously breathing, anesthetized animal. Pressure measurements were repeated three times and the mean value used. Data were collected by Power Lab Data Acquisition system (AD Instruments) and analysed using LabChart 8 software (AD Instruments) by an investigator blinded to the treatment group.

The right lung lobe was harvested and snap frozen in liquid nitrogen or placed in RNAlater^®^ RNA Stabilization Solution for RNA isolation. The left lobe was inflation-fixed (10% formaldehyde in PBS), embedded in paraffin, and sectioned for histology. The heart and liver were collected and snap frozen or placed in RNAlater^®^. Transverse formalin-fixed lung sections were stained with an anti-smooth muscle actin antibody (DAKO M0851) or Verhoeff’s van Gieson stain (EVG) to visualise elastic lamina. Pulmonary vascular remodelling (muscularisation of small intrapulmonary arteries) was determined by counting all muscularised vessels with a diameter smaller than 50 μm in each lung section and expressed as a % of all (muscularised + non-muscularised) vessels. Counting was performed by two observers blinded to treatment.

### *In situ* hybridization

*In situ* hybridization was carried out on paraffin lung sections of untreated mice and Sugen/hypoxia mice (n=3, 5 weeks hypoxia) using miRCURY LNA™ microRNA ISH Optimization kit (Exiqon, cat no 339459). Negative control: LNA™ scrambled microRNA probe, double DIG labelled (40 nM); Positive controls: LNA™ U6snRNA probe, 5’DIG-labeled (1 nM), LNA™ microRNA223 probe, double DIG labelled (40 nM, labels myeloid, granulocytic and monocytic cell lineages in the hematopoietic system). LNA™ microRNA150 probe double DIG labelled (40 nM) was used to study changes in miR-150 levels. Hybridization temperature: 54°C. The sections were incubated with sheep anti-DIG antibody (1:200, Roche Applied Science; cat. no 1333 089), biotinylated donkey anti-sheep antibody (1:200, Sigma, cat no. AP184B), streptavidin-peroxidase conjugate (1:200), followed by DAB/hematoxylin staining. A detailed protocol can be found in miRCURY LNA miRNA Detection Probes Handbook – Qiagen.

### RNAscope^®^

For formalin-fixed, paraffin-embedded lung sections, RNAscope^®^ Multiplex Fluorescent Reagent Kit v2 (Advanced Cell Diagnostics) and TSATM Cyanine 3 & 5, TMR, Fluorescein Evaluation Kit System (PerkinElmer) were used according to manufactures’ protocols(Wang et al., 2012).

Briefly, tissue sections in 5-µm thickness were baked in a dry oven (Agilent G2545A Hybridization Oven, Agilent Technologies) for 1 hour at 60°C, and deparaffinised in xylene, followed by dehydration in 100% ethanol. Tissue sections were then incubated with RNAscope^®^ Hydrogen Peroxide for 10 minutes at room temperature. After washing twice with distilled water, manual target retrieval was performed boiling the sections (100°C to 103°C) in 1X Target Retrieval Reagents using a hot plate for 15 minutes. Slides were then rinsed in deionized water, 100% ethanol, and incubated with RNAscope^®^ Protease Plus at 40°C for 30 minutes in a HybEZ hybridization oven (Advanced Cell Diagnostics, PN 321710/321720). Hybridization with target probes (Mm-Myb-C1, NM_001198914.1; Mm-Notch3-C2, NM_025576.2) was carried out incubating the slides at 40°C for 2 hours. Two different probes/channels (C1-C2) were multiplexed. After washing twice with Wash Buffer, slides were stored overnight in 5x SSC buffer (0.75M NaCl, 0.075M sodium citrate). The following day, the slides were incubated at 40°C with the following reagents: Amplifier 1 (30 min), Amplifier 2 (30 min), Amplifier 3 (15 min); HRP-C1 (15 min), TSA^®^ Plus fluorophore for channel 1 (fluorescein, PerkinElmer; 1:1000; 30 min), HRP blocker (15 min); HRP-C2 (15 min), TSA^®^ Plus fluorophore for channel 2 (cyanine 3, PerkinElmer; 1:1000; 30 min), HRP blocker (15 min). After each hybridization step, slides were washed three times with Wash Buffer at room temperature.

RNAscope hybridisation was combined with immunofluorescence (Sindi et al., 2020, Wang et al., 2012). Tissue was blocked for 1 hour at room temperature with 3% normal horse serum (Vector Laboratories) in 1X PBS containing 0.1% bovine serum albumin (Sigma-Aldrich), and 0.01% sodium azide (Sigma-Aldrich), and then incubated with polyclonal rabbit antibody raised against human von Willebrand Factor (1:500; A0082, Dako), at 4°C overnight. After three washes in PBS, slides were incubated with FITC-labelled Goat Anti-Rabbit antibody (1:100; 111-095-003, Jackson ImmunoResearch Inc.) for 30 minutes at RT. Following immunostaining, tissues were mounted in Vectashield with DAPI and examined under a fluorescent confocal microscope (Leica, TCS SP5, Leica Biosystems, Bretton, Peterborough).

### Cell culture

Human pulmonary artery endothelial cells (HPAECs, Promocell, C-12241) were cultured in endothelial cell growth medium 2 (ECGM2; PromoCell, C-22111) and human pulmonary artery smooth muscle cells (HPASMCs, Lonza, CC-2581) in smooth muscle cell growth medium 2 (SMCGM2, PromoCell, C-22062), as previously described (Wojciak-Stothard et al., 2014). In some experiments, the cells were incubated with human recombinant TNF-α (R&D, 210-TA-020; 10 µg/L), or exposed to hypoxia (5% CO_2_, 2% O_2_) for 18-72 hours.

For non-contact co-culture of HPAECs and HPASMCs, Transwell dishes with 0.4 μm pore polyester membrane inserts (Scientific Laboratory Supplies, UK) were used. HPAECs were seeded into the fibronectin-coated top chambers and cultured in complete ECGM2 medium, whereas HPASMCs were seeded at the bottom of the plate and cultured in complete SMCGM2 for 24 h. The two cell types were then washed with PBS, combined together and co-cultured in endothelial cell basal medium supplemented with 10% FBS (Sigma-Aldrich, F7524), and selected components of ECGM2 supplement pack (PromoCell, C-22111): EGF (2.5 ng/L), FGF (10 ng/L), IGF (20 ng/L) with 1% penicillin and streptomycin.

### Blood-derived human endothelial cells and human lung samples

All investigations were conducted in accordance with the Declaration of Helsinki. Venous blood samples were obtained with the approval of the Brompton Harefield & NHLI and Hammersmith Hospitals Research Ethics committees and informed written consent from healthy volunteers (n=14) and patients with idiopathic PAH (IPAH, n=12). Participants were identified by number. Human endothelial colony forming cells (ECFCs) were derived from peripheral blood samples as previously described (Wojciak-Stothard et al., 2014). Clinical information is shown in Table S2 in the Online Data Supplement.

### Cell Transfection

Briefly, HPAECs were left untreated (control) or were transfected with control miRNA (non-targeting transfection control; Ambion Life Technologies, 4464076) at 20 nmol/L, or miRVana™ has-miR-150-5p, (4464066 Assay ID MC10070;), miRVana™ miRNA inhibitor, (4464084, Assay ID MH10070), both at 20 nmol/L, or control siRNA (non-targeting negative control siRNA; Invitrogen, 4390843) at 10 nmol/L, or siPTPMT1 (4392420 Assay ID s229946) at 10 nmol/L, using Lipofectamine RNAiMAX in antibiotic-free media, following manufacturer’s instruction. After 24 hours, the media were changed and cells were exposed to hypoxia, or treated with TNF-α for 24-72 hours. Alternatively, on the following day, the untransfected and transfected cells were starved for 9 hours before caspase 3/7 assay. Human pcDNA PTPMT1, NM_175732.2 (clone OHu11042; 2B Scientific Ltd. Upper Heyford, UK) was transfected into HPAECs with Lipofectamine RNAiMAX at 2ng/well in a 24-well dish, as recommended by the manuafacturer. Transfection efficiency was measured by the uptake of Cy3™ Dye-Labeled Pre-miR Negative Control (AM17120; Thermo Fisher Scientific) and quantitative real-time PCR (RT-qPCR). All experiments were performed 24 hours after transfection. Transfected cells were exposed to hypoxia (2% O_2_, 5% CO_2_), serum and growth factor depletion or inflammatory cytokines. Cell proliferation and NFκB activity assays were carried out 72 and 48 hours post-transfection, respectively.

### RNA Extraction

RNA was extracted from cultured cells or tissue (∼10 mg) stored in RNALater^®^ using Monarch^®^ Total RNA Miniprep Kit (New England BioLabs). For maximal RNA recovery, tissues was mechanically homogenized using a Kinematica™ Polytron™ PT 1300 D and incubated at 55°C for 5 minutes with Proteinase K following manufacturer’s instructions. To remove any residual DNA that may affect downstream applications, an On-Column DNase I digestion was also performed. RNA concentration and purity was evaluated using NanoDrop 2000 spectrophotometer (Thermo Scientific). The A260/230 and A260/280 ratios were used to assess the presence of contaminants. RNA was then stored at −80°C for later experiments.

### Real-time quantitative PCR

Input RNA (50-100 ng/µL) was reverse-transcribed using LunaScript^®^ RT SuperMix Kit (New England BioLabs) or TaqMan MicroRNA Reverse Transcription Kit (Thermo Fisher Scientific) and custom Multiplex RT Primer pool in a SimpliAmp™ Thermal Cycler (Applied Biosystems), according to the manufacturer’s instructions. The multiplex RT primer pool consisted of primers for miR-150-5p and U6 (Thermo Fisher Scientific). No-template samples where included as negative controls.

TaqMan^®^ miRNA Assays for hsa-miR-150-5p (Assay ID 000473), and U6 snRNA (Assay ID 001973), and TaqMan® Gene Expression Assays for PTPMT1 (Hs00378514_m1, Mm00458631_m1), SERPINE1 (Hs00167155_m1), PERP (Hs00953482_g1), DUSP5 (Hs00244839_m1), c-MYB (Hs00920556_m1, Mm00501741_m1), NOTCH3 (Hs01128537_m1, Mm01345646_m1), Col1a1 (Mm00801666_g1), Rcan1 (Mm01213406_m1), Tgfb1 (Mm01178820_m1), and GAPDH (Hs02786624_g1, Mm99999915_g1; all Thermo Fisher Scientific), were used to perform quantitative PCR (qPCR). No-template samples were included as negative controls and all PCRs were performed in triplicate. The reaction was performed on a QuantStudio 12K Flex Real-Time PCR System (Applied Biosystems). Data were analysed using QuantStudio 12K Flex Software version 1.2 (Applied Biosystems). For relative quantification, the data were analysed using the 2^−ΔΔCt^ method, where U6 snRNA and GAPDH were used as endogenous normalization controls for miR-150 and gene expression, respectively.

### RNA-Sequencing and identification of signaling mediators of miR-150

10µl of RNA (250-300 ng/µl) extracted from cells transfected with miR-150 mimic or scrambled control in three independent experiments, were sent in to Imperial BRC Genomics Facility (Imperial College of London, UK) for next-generation RNA-sequencing RNA quality and quantity were assessed using an Agilent 2100 Bioanalyzer (Agilent Technologies) and a Qubit 4 Fluorometer (Thermo Fisher Scientific). RNA libraries were prepared using TruSeq^®^ Stranded mRNA HT Sample Prep Kit (Illumina Inc., USA) according to the manufacturer’s protocol as previously described(Sindi et al., 2020). Libraries were run over 4 lanes (2 × 100 bp) on a HiSeq 2500 (Illumina Inc.) resulting in an average of 34.4 million reads per sample. Sequence data was de-multiplexed using bcl2fastq2 Conversion Software v2.18 (Illumina Inc.) and quality analysed using FastQC (http://www.bioinformatics.babraham.ac.uk/projects/fastqc). Transcripts from paired-end stranded RNA-Seq data were quantified with Salmon v 0.8.2 using hg38 reference transcripts (Love, Huber et al., 2014, Patro, Duggal et al., 2017). Count data was normalised to accommodate known batch effects and library size using DESeq2 (Anders, Pyl et al., 2015). Pairwise differential expression analysis was performed based on a model using the negative binomial distribution and p-values were adjusted for multiple test correction using the Benjamini-Hochberg procedure (Hochberg & Benjamini, 1990). Genes were considered differentially expressed if the adjusted p-value was greater than 0.05 and there was at least a 1.5 fold change in expression. miRNA target prediction was carried out with TargetScan Human, miRecords and Ingenuity Expert Findings. Gene enrichment was carried out using Ingenuity Pathway Analysis (IPA, Qiagen).

### Caspase 3/7 apoptosis assay

Cells were incubated in serum- and growth factor-depleted medium for 9 hours to induce apoptosis. Apoptosis was measured using Cell Meter™ Caspase 3/7 Activity Apoptosis Assay Kit (AAT Bioquest, ABD-22796). Fluorescence intensity was analysed in Glomax™ luminometer at Ex/Em = 490/525 nm.

### NFκB luciferase reporter assay

NFκB activity was measured in luciferase reporter assay (Wojciak-Stothard et al., 2014) in the Glomax™ luminometer.

### EdU Proliferation Assay

The EdU Cell Proliferation Assay Kit (EdU-594, EMD Millipore Corp, USA, 17-10527) was used to measure cell proliferation, according to the manufacturer’s protocol. Proliferation of ECFCs from healthy individuals and IPAH patients was evaluated in serum-reduced, growth factor-depleted media.

### Seahorse Bioenergetics Assay

Oxygen consumption (OCR) and extracellular acidification rates (ECAR) were measured in Seahorse Extracellular Flux Analyzer using XF24 (Seahorse Bioscience, North Billerica, MA) and Seahorse XF Mito Stress Test Kit (Agilent, 103015-100). 4 × 10^4^ cells were plated into each well prior to the assay. Cells cultured on the Seahorse XF Cell Culture microplates were left untreated or were transfected with miR-150 or PTPMT1, as previously described for 24h overexpression. The sensor cartridge was hydrated at 37°C in Seahorse XF Calibrant overnight in a non-CO_2_ incubator.

The assay medium was prepared by supplementing Seahorse XF Base Medium with 1mM pyruvate (S8636), 2mM glutamate and glucose (G8540) and 10mM glucose (G8769), warming it up to 37°C and adjusting pH to 7.4. All compounds were warmed up to room temperature. 1μM oligomycin, 1μM FCCP and 0.5μM rotenone/antimycin A provided with the kit were loaded into the appropriate ports of hydrated sensor cartridge. The cells were incubated with assay medium for 1h before Seahorse XF Mito Stress Test. OCR and ECAR were normalized to the protein concentration.

### Immunostaining

Immunostaining of paraffin embedded lung sections was carried out as previously described (Wojciak-Stothard et al., 2014).

To stain mitochondria, HPAECs cultured on Thermanox® Plastic Coverslips (13 mm) were washed twice in PBS, fixed in 4% formaldehyde for 15 min at room temperature, washed in PBS and permeabilised with 0.1% Triton X-100 (Sigma-Aldrich, 234729) in PBS for 10 min. The cells were then rinsed with PBS, blocked in 10% normal goat serum (Vector Laboratories, S-1000) for 1 h and incubated with mouse anti-mitochondria antibodies (Abcam, ab92824) diluted 1:100 in PBS in 5% BSA in a humidified chamber overnight. Cells were then rinsed 3 times with PBS and incubated with FITC-Goat Anti-Mouse IgG (Jackson ImmunoResearch Inc.,115-095-003; 1:200) with tetramethylrhodamine (TRITC)-phalloidin (1 μg/mL; Sigma-Aldrich, UK, P1951) for 1h. Following immunostaining, cells were mounted in Vectashield Antifade Mounting Medium containing nuclear stain DAPI (Vector Laboratories, H-1200) and examined under the fluorescent confocal microscope (Leica, TCS SP5, Leica Biosystems, Bretton, Peterborough).

### Cardiolipin measurement

Quantification of cardiolipin in cells and tissues was carried out with Cardiolipin Assay Kit (BioVision, cat. K944-100), according to the manufacturer’s instructions.

### Mitochondrial fragmentation count and mitochondrial content

Mitochondrial were immunolabelled, as described above. Mitochondrial fragmentation (area taken by mitochondrial particles < 2μm in length) (Farrand, Kim et al., 2013) and total mitochondrial coverage (area taken by all mitochondria) were determined using NIP2 image software (Martinez & Cupitt, 2005). The 2μm cut-off size was optimal(Farrand et al., 2013) in selection of mitochondria unassociated with mitochondrial network. Briefly, the acquired images were filtered (median), thresholded, and binarized to identify individual mitochondrial segments and score the total area of fragmented mitochondria. This value was normalized to the total mitochondrial area (in pixels) in each cell, to define the individual cell’s MFC. For each intervention 20 randomly selected cells were analysed in 3 separate experiments (Hong, Kutty et al., 2013).

### Statistics

All experiments were performed at least in triplicate and measurements were taken from distinct samples. Data are presented as mean±SEM. Normality of data distribution was assessed with Shapiro-Wilk test in GraphPad Prism 7.03. Comparisons between 2 groups were made with Student t test or Mann-Whitney’s U test, whereas ≥3 groups were compared by use of ANOVA with Tukey’s post hoc test or Kruskal-Wallis with Dunn’s post hoc test, as appropriate. Statistical significance was accepted at p<0.05.

## ACKNOWLEDGEMENTS

This research was supported by PhD Studentships from the Government of Saudi Arabia (Mai Alzaydi) and British Heart Foundation project grant PG/16/4/31849. We thank the staff of the Imperial NIHR/Imperial Clinical Research Facility, Hammersmith Hospital (London UK) and Dr John Wharton (NHLI) for their help in acquiring cells from IPAH patients. We also thank Prof. Anna Randi and Dr Graeme Birdsay (NHLI, Imperial College London) for providing Cdh5(PAC)-iCreERT2 mice and sharing experimental protocols used to obtain the endothelium-specific mir-150 knockout mice

## AUTHOR CONTRIBUTIONS

GR, KBJ, VAS, MA, CCM and BWS performed *in vitro* experiments, analysed data; KBJ, GR and VBA-S in vitro and in vivo experiments, immunohistochemistry, analysed data; MA analysis of mitochondrial fragmentation; MRW critical analysed the manuscript; CCM analysed RNA-Seq data; MRW provided IPAH patient data; BWS secured funding, performed experiments, wrote the manuscript.

## THE PAPER EXPLAINED

### Problem

Reduced circulating levels of miR-150 correlate with poor survival in pulmonary arterial hypertension. Links between endothelial miR-150 and vascular dysfunction in PAH are not well understood.

### Results

Lung endothelium-targeted miR-150 delivery prevents the development of the disease, while endothelial knockdown of miR-150 has adverse effects. Beneficial effects of miR-150 *in vitro* and *in vivo* are linked with PTEN-like mitochondrial phosphatase (PTPMT1)-dependent biosynthesis of mitochondrial phospholipid cardiolipin and reduced expression of pro-apoptotic, pro-inflammatory and pro-fibrotic genes.

### Impact

Improvement of mitochondrial function by miR-150-PTPMT1-cardiolipin signaling is likely to facilitate adaptation of lung and heart to high energy demand created by mechanical workload, inflammation and hypoxia in stress conditions.

## DATA AVAILABILITY

Accession numbers for RNA sequencing data will be provided upon provisional acceptance of the manuscript

## References

Abdul-Salam VB, Russomanno G, Chien-Nien C, Mahomed AS, Yates LA, Wilkins MR, Zhao L, Gierula M, Dubois O, Schaeper U, Endruschat J, Wojciak-Stothard B (2019) CLIC4/Arf6 Pathway. Circ Res 124: 52–65

Anders S, Pyl PT, Huber W (2015) HTSeq--a Python framework to work with high-throughput sequencing data. Bioinformatics 31: 166–9

Archer SL (2017) Pyruvate Kinase and Warburg Metabolism in Pulmonary Arterial Hypertension: Uncoupled Glycolysis and the Cancer-Like Phenotype of Pulmonary Arterial Hypertension. Circulation 136: 2486–2490

Bertero T, Handen AL, Chan SY (2018) Factors Associated with Heritable Pulmonary Arterial Hypertension Exert Convergent Actions on the miR-130/301-Vascular Matrix Feedback Loop. Int J Mol Sci 19

Birdsey GM, Shah AV, Dufton N, Reynolds LE, Osuna Almagro L, Yang Y, Aspalter IM, Khan ST, Mason JC, Dejana E, Gottgens B, Hodivala-Dilke K, Gerhardt H, Adams RH, Randi AM (2015) The endothelial transcription factor ERG promotes vascular stability and growth through Wnt/beta-catenin signaling. Dev Cell 32: 82–96

Boucher J, Gridley T, Liaw L (2012) Molecular pathways of notch signaling in vascular smooth muscle cells. Front Physiol 3: 81

Cao M, Hou D, Liang H, Gong F, Wang Y, Yan X, Jiang X, Wang C, Zhang J, Zen K, Zhang CY, Chen X (2014) miR-150 promotes the proliferation and migration of lung cancer cells by targeting SRC kinase signalling inhibitor 1. Eur J Cancer 50: 1013–24

Caruso P, Dunmore BJ, Schlosser K, Schoors S, Dos Santos C, Perez-Iratxeta C, Lavoie JR, Zhang H, Long L, Flockton AR, Frid MG, Upton PD, D’Alessandro A, Hadinnapola C, Kiskin FN, Taha M, Hurst LA, Ormiston ML, Hata A, Stenmark KR et al. (2017) Identification of MicroRNA-124 as a Major Regulator of Enhanced Endothelial Cell Glycolysis in Pulmonary Arterial Hypertension via PTBP1 (Polypyrimidine Tract Binding Protein) and Pyruvate Kinase M2. Circulation 136: 2451–2467

Chandra SM, Razavi H, Kim J, Agrawal R, Kundu RK, de Jesus Perez V, Zamanian RT, Quertermous T, Chun HJ (2011) Disruption of the apelin-APJ system worsens hypoxia-induced pulmonary hypertension. Arterioscler Thromb Vasc Biol 31: 814–20

Chen LL, Zmuda EJ, Talavera MM, Frick J, Brock GN, Liu Y, Klebanoff MA, Trittmann JK (2020) Dual-specificity phosphatase (DUSP) genetic variants predict pulmonary hypertension in patients with bronchopulmonary dysplasia. Pediatr Res 87: 81–87

Chen M, Shen C, Zhang Y, Shu H (2017) MicroRNA-150 attenuates hypoxia-induced excessive proliferation and migration of pulmonary arterial smooth muscle cells through reducing HIF-1alpha expression. Biomed Pharmacother 93: 861–868

Cheng J, Nanayakkara G, Shao Y, Cueto R, Wang L, Yang WY, Tian Y, Wang H, Yang X (2017) Mitochondrial Proton Leak Plays a Critical Role in Pathogenesis of Cardiovascular Diseases. Adv Exp Med Biol 982: 359–370

Choi SY, Gonzalvez F, Jenkins GM, Slomianny C, Chretien D, Arnoult D, Petit PX, Frohman MA (2007) Cardiolipin deficiency releases cytochrome c from the inner mitochondrial membrane and accelerates stimuli-elicited apoptosis. Cell Death Differ 14: 597–606

Culley MK, Chan SY (2018) Mitochondrial metabolism in pulmonary hypertension: beyond mountains there are mountains. J Clin Invest 128: 3704–3715

Dasgupta A, Wu D, Tian L, Xiong PY, Dunham-Snary KJ, Chen KH, Alizadeh E, Motamed M, Potus F, Hindmarch CCT, Archer SL (2020) Mitochondria in the Pulmonary Vasculature in Health and Disease: Oxygen-Sensing, Metabolism, and Dynamics. Compr Physiol 10: 713–765

Deng P, Chen L, Liu Z, Ye P, Wang S, Wu J, Yao Y, Sun Y, Huang X, Ren L, Zhang A, Wang K, Wu C, Yue Z, Xu X, Chen M (2016) MicroRNA-150 Inhibits the Activation of Cardiac Fibroblasts by Regulating c-Myb. Cell Physiol Biochem 38: 2103–22

Desjarlais M, Dussault S, Dhahri W, Mathieu R, Rivard A (2017) MicroRNA-150 Modulates Ischemia-Induced Neovascularization in Atherosclerotic Conditions. Arterioscler Thromb Vasc Biol 37: 900–908

Dudek J (2017) Role of Cardiolipin in Mitochondrial Signaling Pathways. Front Cell Dev Biol 5: 90

Duong HT, Comhair SA, Aldred MA, Mavrakis L, Savasky BM, Erzurum SC, Asosingh K (2011) Pulmonary artery endothelium resident endothelial colony-forming cells in pulmonary arterial hypertension. Pulm Circ 1: 475–86

Eichstaedt CA, Song J, Viales RR, Pan Z, Benjamin N, Fischer C, Hoeper MM, Ulrich S, Hinderhofer K, Grunig E (2017) First identification of Kruppel-like factor 2 mutation in heritable pulmonary arterial hypertension. Clin Sci (Lond) 131: 689–698

Farrand L, Kim JY, Im-Aram A, Suh JY, Lee HJ, Tsang BK (2013) An improved quantitative approach for the assessment of mitochondrial fragmentation in chemoresistant ovarian cancer cells. PLoS One 8: e74008

Fehring V, Schaeper U, Ahrens K, Santel A, Keil O, Eisermann M, Giese K, Kaufmann J (2014) Delivery of therapeutic siRNA to the lung endothelium via novel Lipoplex formulation DACC. Mol Ther 22: 811–20

Freund-Michel V, Khoyrattee N, Savineau JP, Muller B, Guibert C (2014) Mitochondria: roles in pulmonary hypertension. Int J Biochem Cell Biol 55: 93–7

Ghose J, Bhattacharyya NP (2015) Transcriptional regulation of microRNA-100, -146a, and -150 genes by p53 and NFkappaB p65/RelA in mouse striatal STHdh(Q7)/ Hdh(Q7) cells and human cervical carcinoma HeLa cells. RNA Biol 12: 457–77

Gubrij IB, Pangle AK, Pang L, Johnson LG (2016) Reversal of MicroRNA Dysregulation in an Animal Model of Pulmonary Hypertension. PLoS One 11: e0147827

Havrda MC, Johnson MJ, O’Neill CF, Liaw L (2006) A novel mechanism of transcriptional repression of p27kip1 through Notch/HRT2 signaling in vascular smooth muscle cells. Thromb Haemost 96: 361–70

Hergenreider E, Heydt S, Treguer K, Boettger T, Horrevoets AJ, Zeiher AM, Scheffer MP, Frangakis AS, Yin X, Mayr M, Braun T, Urbich C, Boon RA, Dimmeler S (2012) Atheroprotective communication between endothelial cells and smooth muscle cells through miRNAs. Nat Cell Biol 14: 249–56

Hochberg Y, Benjamini Y (1990) More powerful procedures for multiple significance testing. Stat Med 9: 811–8

Hong Z, Kutty S, Toth PT, Marsboom G, Hammel JM, Chamberlain C, Ryan JJ, Zhang HJ, Sharp WW, Morrow E, Trivedi K, Weir EK, Archer SL (2013) Role of dynamin-related protein 1 (Drp1)-mediated mitochondrial fission in oxygen sensing and constriction of the ductus arteriosus. Circ Res 112: 802–15

Li J, Zhang Y, Liu Y, Dai X, Li W, Cai X, Yin Y, Wang Q, Xue Y, Wang C, Li D, Hou D, Jiang X, Zhang J, Zen K, Chen X, Zhang CY (2013) Microvesicle-mediated transfer of microRNA-150 from monocytes to endothelial cells promotes angiogenesis. J Biol Chem 288: 23586–96

Li X, Zhang X, Leathers R, Makino A, Huang C, Parsa P, Macias J, Yuan JX, Jamieson SW, Thistlethwaite PA (2009) Notch3 signaling promotes the development of pulmonary arterial hypertension. Nat Med 15: 1289–97

Lim SL, Lam CS, Segers VF, Brutsaert DL, De Keulenaer GW (2015) Cardiac endothelium-myocyte interaction: clinical opportunities for new heart failure therapies regardless of ejection fraction. Eur Heart J 36: 2050–2060

Liu F, Di Wang X (2019) miR-150-5p represses TP53 tumor suppressor gene to promote proliferation of colon adenocarcinoma. Sci Rep 9: 6740

Love MI, Huber W, Anders S (2014) Moderated estimation of fold change and dispersion for RNA-seq data with DESeq2. Genome Biol 15: 550

Martinez K, Cupitt J (2005) VIPS – a highly tuned image processing software architecture. Ieee Image Proc: 2485–2488

Mason EF, Rathmell JC (2011) Cell metabolism: an essential link between cell growth and apoptosis. Biochim Biophys Acta 1813: 645–54

Monticelli S, Ansel KM, Xiao C, Socci ND, Krichevsky AM, Thai TH, Rajewsky N, Marks DS, Sander C, Rajewsky K, Rao A, Kosik KS (2005) MicroRNA profiling of the murine hematopoietic system. Genome Biol 6: R71

Negi V, Chan SY (2017) Discerning functional hierarchies of microRNAs in pulmonary hypertension. JCI Insight 2: e91327

Paradies G, Paradies V, Ruggiero FM, Petrosillo G (2019) Role of Cardiolipin in Mitochondrial Function and Dynamics in Health and Disease: Molecular and Pharmacological Aspects. Cells 8

Patro R, Duggal G, Love MI, Irizarry RA, Kingsford C (2017) Salmon provides fast and bias-aware quantification of transcript expression. Nat Methods 14: 417–419

Paulin R, Michelakis ED (2014) The metabolic theory of pulmonary arterial hypertension. Circ Res 115: 148–64

Pitulescu ME, Schmidt I, Benedito R, Adams RH (2010) Inducible gene targeting in the neonatal vasculature and analysis of retinal angiogenesis in mice. Nat Protoc 5: 1518–34

Rawat DK, Alzoubi A, Gupte R, Chettimada S, Watanabe M, Kahn AG, Okada T, McMurtry IF, Gupte SA (2014) Increased reactive oxygen species, metabolic maladaptation, and autophagy contribute to pulmonary arterial hypertension-induced ventricular hypertrophy and diastolic heart failure. Hypertension 64: 1266–74

Rhodes CJ, Wharton J, Boon RA, Roexe T, Tsang H, Wojciak-Stothard B, Chakrabarti A, Howard LS, Gibbs JS, Lawrie A, Condliffe R, Elliot CA, Kiely DG, Huson L, Ghofrani HA, Tiede H, Schermuly R, Zeiher AM, Dimmeler S, Wilkins MR (2013) Reduced microRNA-150 is associated with poor survival in pulmonary arterial hypertension. Am J Respir Crit Care Med 187: 294–302

Ryan JJ, Archer SL (2014) The right ventricle in pulmonary arterial hypertension: disorders of metabolism, angiogenesis and adrenergic signaling in right ventricular failure. Circ Res 115: 176–88

Saini-Chohan HK, Dakshinamurti S, Taylor WA, Shen GX, Murphy R, Sparagna GC, Hatch GM (2011) Persistent pulmonary hypertension results in reduced tetralinoleoyl-cardiolipin and mitochondrial complex II + III during the development of right ventricular hypertrophy in the neonatal pig heart. Am J Physiol Heart Circ Physiol 301: H1415–24

Schermuly RT, Ghofrani HA, Wilkins MR, Grimminger F (2011) Mechanisms of disease: pulmonary arterial hypertension. Nat Rev Cardiol 8: 443–55

Shen J, Liu X, Yu WM, Liu J, Nibbelink MG, Guo C, Finkel T, Qu CK (2011) A critical role of mitochondrial phosphatase Ptpmt1 in embryogenesis reveals a mitochondrial metabolic stress-induced differentiation checkpoint in embryonic stem cells. Mol Cell Biol 31: 4902–16

Simone TM, Higgins CE, Czekay RP, Law BK, Higgins SP, Archambeault J, Kutz SM, Higgins PJ (2014) SERPINE1: A Molecular Switch in the Proliferation-Migration Dichotomy in Wound-“Activated” Keratinocytes. Adv Wound Care (New Rochelle) 3: 281–290

Sindi HA, Russomanno G, Satta S, Abdul-Salam VB, Jo KB, Qazi-Chaudhry B, Ainscough AJ, Szulcek R, Jan Bogaard H, Morgan CC, Pullamsetti SS, Alzaydi MM, Rhodes CJ, Piva R, Eichstaedt CA, Grunig E, Wilkins MR, Wojciak-Stothard B (2020) Therapeutic potential of KLF2-induced exosomal microRNAs in pulmonary hypertension. Nat Commun 11: 1185

Vaschetto LM (2018) miRNA activation is an endogenous gene expression pathway. RNA Biol 15: 826–828

Voelkel NF, Quaife RA, Leinwand LA, Barst RJ, McGoon MD, Meldrum DR, Dupuis J, Long CS, Rubin LJ, Smart FW, Suzuki YJ, Gladwin M, Denholm EM, Gail DB, National Heart L, Blood Institute Working Group on C, Molecular Mechanisms of Right Heart F (2006) Right ventricular function and failure: report of a National Heart, Lung, and Blood Institute working group on cellular and molecular mechanisms of right heart failure. Circulation 114: 1883–91

Wang F, Flanagan J, Su N, Wang LC, Bui S, Nielson A, Wu X, Vo HT, Ma XJ, Luo Y (2012) RNAscope: a novel in situ RNA analysis platform for formalin-fixed, paraffin-embedded tissues. Mol Diagn 14: 22–9

Wang W, Li C, Li W, Kong L, Qian A, Hu N, Meng Q, Li X (2014) MiR-150 enhances the motility of EPCs in vitro and promotes EPCs homing and thrombus resolving in vivo. Thromb Res 133: 590–8

Wang Y, Nakayama M, Pitulescu ME, Schmidt TS, Bochenek ML, Sakakibara A, Adams S, Davy A, Deutsch U, Luthi U, Barberis A, Benjamin LE, Makinen T, Nobes CD, Adams RH (2010) Ephrin-B2 controls VEGF-induced angiogenesis and lymphangiogenesis. Nature 465: 483–6

Wang Z, Yang K, Zheng Q, Zhang C, Tang H, Babicheva A, Jiang Q, Li M, Chen Y, Carr SG, Wu K, Zhang Q, Balistrieri A, Wang C, Song S, Ayon RJ, Desai AA, Black SM, Garcia JGN, Makino A et al. (2019) Divergent changes of p53 in pulmonary arterial endothelial and smooth muscle cells involved in the development of pulmonary hypertension. Am J Physiol Lung Cell Mol Physiol 316: L216–L228

Wojciak-Stothard B, Abdul-Salam VB, Lao KH, Tsang H, Irwin DC, Lisk C, Loomis Z, Stenmark KR, Edwards JC, Yuspa SH, Howard LS, Edwards RJ, Rhodes CJ, Gibbs JS, Wharton J, Zhao L, Wilkins MR (2014) Aberrant chloride intracellular channel 4 expression contributes to endothelial dysfunction in pulmonary arterial hypertension. Circulation 129: 1770–80

Wu Q, Jin H, Yang Z, Luo G, Lu Y, Li K, Ren G, Su T, Pan Y, Feng B, Xue Z, Wang X, Fan D (2010) MiR-150 promotes gastric cancer proliferation by negatively regulating the pro-apoptotic gene EGR2. Biochem Biophys Res Commun 392: 340–5

You XM, Mungrue IN, Kalair W, Afroze T, Ravi B, Sadi AM, Gros R, Husain M (2003) Conditional expression of a dominant-negative c-Myb in vascular smooth muscle cells inhibits arterial remodeling after injury. Circ Res 92: 314–21

Zhang J, Guan Z, Murphy AN, Wiley SE, Perkins GA, Worby CA, Engel JL, Heacock P, Nguyen OK, Wang JH, Raetz CR, Dowhan W, Dixon JE (2011) Mitochondrial phosphatase PTPMT1 is essential for cardiolipin biosynthesis. Cell Metab 13: 690–700

Zhao M, Chen N, Li X, Lin L (2019) MiR-629 regulates hypoxic pulmonary vascular remodelling by targeting FOXO3 and PERP. J Cell Mol Med 23: 5165–5175

